# IQGAP2 acts as a tumor suppressor in breast cancer and its reduced expression promotes cancer growth and metastasis by MEK/ERK signalling pathways

**DOI:** 10.1101/651034

**Authors:** Dinesh Kumar, Md. Khurshidul Hassan, Niharika Pattanaik, Nachiketa Mohapatra, Manjusha Dixit

**Author notes:** Corresponding Author: (MD).

## Abstract

IQGAP2 is a member of IQGAPs scaffolding protein family. It has been reported as a tumor suppressor in various cancers, as well as, an oncogene in some cancers, suggesting organ specific role. Need to identify therapeutic targets which function in ER/PR independent way, prompted us to explore role of IQGAP2 in molecular mechanism in breast cancer, which was completely unknown. In vitro studies in estrogen receptor positive breast cancer cell line (MCF7) showed that low IQGAP2 expression results in increased cell proliferation, migration and invasion of cells whereas an opposite effect was observed with ectopic expression of IQGAP2. Triple negative breast cancer cell line (MDA-MB-468), with IQGAP2 depletion showed similar effect, supporting its role in ER/PR independent manner. Furthermore, we found that reduced IQGAP2 expression induces the expression of EMT markers; twist and N-cadherin and decreases the expression of MET marker, E-cadherin via the MEK/ERK pathway but not via AKT pathway. Validation of findings in patients showed a reduced IQGAP2 expression in breast cancer tissues compared to normal tissue. Patients with low levels of IQGAP2 showed correlation with higher tumor stage. Our results suggest that IQGAP2 acts as a tumor suppressor and its down regulation results in cell growth, cell invasion and EMT through the MEK/ERK signalling pathways and it hence may be a potential therapeutic target in breast cancer.

## Introduction

Breast cancer is at the top of all cancer incidences in women all around the world. According to SEER, 268,600 new cases of female breast cancer is estimated by the end of 2019 in US population [1]. Despite the advancement in breast tumor therapies, this is the second highest cause of cancer deaths, in women of developed countries [2–4]. Although the survival rate of breast cancer patients has improved, long-term survival still remains low due to drug resistance and relapse of metastasis [5]. Many genes have been recognized which have role in initiation or progression of breast cancer [4, 6–8] but lacunae in understanding of pathogenesis leaves a lot of scope to identify additional players, especially the one which can work independent of breast cancer molecular subtype.

IQGAPs (IQ Motif Containing GTPase Activating Proteins) are a class of scaffolding proteins and are evolutionarily conserved from lower to the higher eukaryotes [9]. Three members of this family namely, IQGAP1, IQGAP2 and IQGAP3, share high similarity at five conserved domains [10]. Despite the presence of similar domains, they show diverse cellular functions [9]. IQGAP1 and IQGAP3 have been found to be upregulated in many cancers and promote tumor growth and metastasis [11–15], whereas the role of IQGAP2 is not fully understood in cancers. Most of the reports showed a reduced expression of IQGAP2 in HCC, prostate, gastric and ovary cancer [16–19], suggesting it to be a tumor suppressor but in couple of studies it was found to be an oncogene too [20, 21]. Our previous work also showed reduced expression level of IQGAP2 in breast cancer, using Oncomine data analysis [22]. Based upon these observations we hypothesised that IQGAP2 might be implicated in breast cancer tumorigenesis as a tumor suppressor gene.

In this study, to find out if the role of IQGAP2 in breast cancer tumorigenesis is independent of molecular subtype, we first explored the effect of IQGAP2 expression level, on tumorigenic properties of ER/PR positive breast cancer cell line, followed by similar study in triple negative breast cancer cell line. We also explored the precise mechanism/pathway by which IQGAP2 regulates invasion and EMT events in vitro. Our findings provide a potential therapeutic target to inhibit breast cancer progression and metastasis.

## Materials and Methods

### Breast cancer patient sample collection

The archival, formalin-fixed paraffin-embedded (FFPE) tissue blocks of 95 breast cancer patients were collected from the SRL Diagnostic Lab, Bhubaneswar and Department of Pathology, Apollo hospital, Bhubaneswar. The study was approved by the Institutional Ethics Committee, NISER, Bhubaneswar (protocol no. NISER/IEC/2016-01). The clinicopathological characteristics of each patient were recorded, which included age, histological type, tumor size, Tumor-Node-Metastasis (TNM) stage and lymphovascular invasion.

### Immunohistochemistry

FFPE block of cancer and matched normal tissue of each patient, was cut into 5 µm thin sections and, placed on poly L-lysine coated glass slides. The sections were deparaffinized by heating slides at 80°C for 1 hour followed by rehydration as follows: Xylene- 3 min (twice), 100% ethanol- 3 min, 90% ethanol- 3 min, 70% ethanol- 3 min, 50% ethanol- 3 min, water- 2 min, and water- 2 minutes. The epitope retrieval was performed, in citrate low pH (pH 6.8) retrieval buffer, by heat induction. Blocking was done in Envision Peroxidase Blocker (Dako) for 15 minutes followed by incubation with 1:100 diluted IQGAP2 primary antibody (Abcam), for 1 hour at room temperature. Subsequently, the sections were incubated with Envision Flex HRP secondary antibody (Dako) for 30 minutes and color was developed using liquid DAB Substrate. Finally, the sections were counterstained with hematoxylin. Two pathologists (NP and NM) did reporting independently, to nullify biasness.

The reporting of IQGAP2 IHC data was done as per the Allred scoring system [23]. In brief, Allred score was calculated by adding the values of intensity score (0 for negative, 1 for weak, 2 for moderate and, 3 for strong) and proportion score (0 – no cells are IQGAP2 +ve, 1 – ≤1% of cells are IQGAP2 +ve, 2 – 1%–10% of cells are IQGAP2 +ve, 3 – 11%–33% of cells are IQGAP2 +ve, 4 – 34%–66% of cells are IQGAP2 +ve, 5 – 67%–100% of cells are IQGAP2 +ve) of cells. For statistical analyses, the Allred scores of 0-2 were treated as negative or weak, 3-6 as moderate and, 7-8 as intense/strong expression levels. For correlation study among different histopathological parameters and IQGAP2 IHC score, Allred score were divided into two groups, scores of 0-2 were considered negative and scores of 3- 8 were considered positive.

### IHC in human breast cancer tissue arrays

We also determined the expression of IQGAP2 in human breast cancer tissue microarray (US BioMax, BC081120c). In this array, 5 µm sections of 100 breast cancer tumor tissues and 10 normal adjacent tissues of breast were available. The immunohistochemistry and scoring of this array was performed in same way as described in immunohistochemistry section. Histopathological and clinical information of the patient i.e. age, histological grade, lymphnode status, TNM staging were supplied with the array.

### Cell culture, plasmids and stable line preparation

The breast cancer cell lines, MCF7 and MDA-MB-468 were purchased from cell repository, NCCS, Pune, India. MCF7 were grown and maintained in DMEM (HiMedia) supplemented with 10% FBS (US origin, HiMedia), ampicillin (HiMedia, 100 IU/ml), streptomycin (HiMedia, 100 µg/ml) whereas MDA-MB-468 were maintained in RPMI (HiMedia) supplemented with 10% FBS (US origin, HiMedia), ampicillin (HiMedia, 100 IU/ml), streptomycin (HiMedia, 100 µg/ml). The cells were cultured at 37°C in 5% CO2. For ectopic IQGAP2 expression and knockdown, IQGAP2 expression vector (pCMV6_IQGAP2_myc), IQGAP2 knockdown vector (pLKO.1_IQGAP2) and their respective control vectors were purchased from OriGene and Sigma, respectively. For transient expression or knockdown of IQGAP2, 1×10^6^ cells were transfected with 1:2 ratio of plasmid and Lipofectamine 3000 (Thermo Scientific). The protein or mRNA were isolated after 36-48 hours of transfection. For stable expression cell line preparation, IQGAP2 expression vector or knockdown vectors were transfected in MCF7 or MDA-MB-468 and grown in 10% DMEM or RPMI media supplemented with G418 (1000 µg/ml) or puromycin (1µg/ml) antibiotics for 10-15 days till the isolated colonies appeared. Few isolated colonies were grown in individual wells and positive stable colonies of IQGAP2 were screened by western blot. Next, the positive colonies were expanded and used for all in-vitro or in-vivo studies. Hereafter these cells will be referred as MCF7 IQGAP2_Ex (MCF7 with IQGAP2 ectopic expression), MCF7 Control_EV (MCF7 with empty expression vector), MCF7 IQGAP2_KD (MCF7 with IQGAP2 knock down), MCF7 Control_Sc (MCF7 having knock down vector with scrambled sequence), MDA-MB-468 IQGAP2_KD (MDA-MB-468 with IQGAP2 knock down), MDA-MB-468 Control_Sc (MDA-MB-468 having knock down vector with scrambled sequence).

### Quantitative real-time PCR

To quantify the endogenous RNA level of IQGAP2, RNeasy Mini Kit (Qiagen) was used to isolate total RNA from different breast cancer cell lines i.e. MCF7, T-47D, MDA-MB-453, MDA-MB-468, MCF 10A and MDA-MB-231 as per the manufacturer’s protocol. cDNA was synthesised using Superscript IV Reverse Transcriptase cDNA synthesis kit (Invitrogen). The primer sequences for genes screened in this study have been shown in the table 1. Quantitative real time PCR was performed, using the Fast Start Universal SYBR Green Master Mix (Roche). ABI 7500 platform (Applied Biosystems) was used to perform the reaction. The experiment was conducted using three biological and three technical replicates. We used GAPDH as an internal control. Relative expression was calculated using the 2^−ΔΔCT^ method.

**Table 1.**
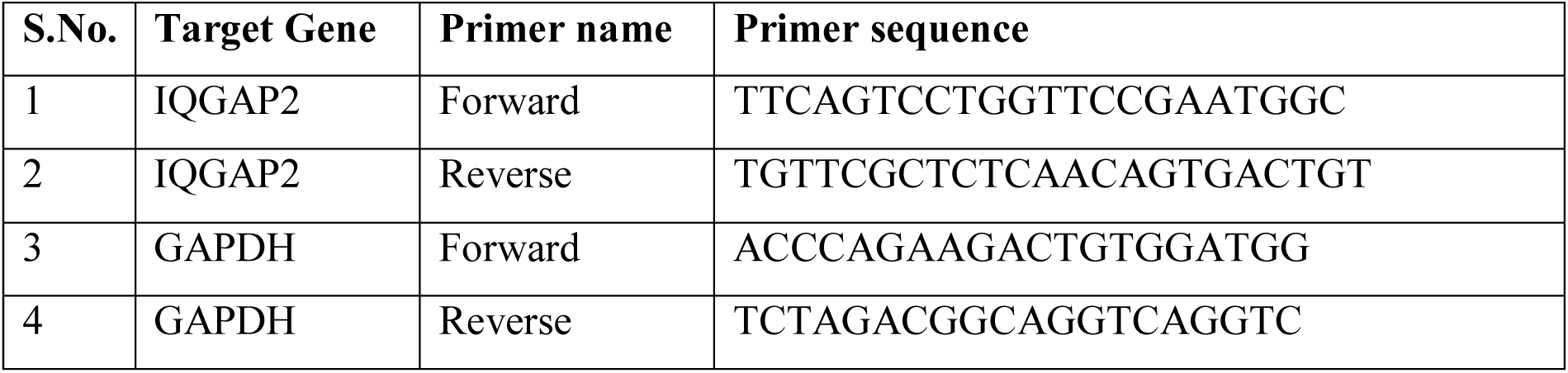
List of primers used in qRT-PCR.

### Western blot analysis

Cells were lysed in ice-cold RIPA lysis buffer (Pierce), with a mixture of protease and phosphatase inhibitor (Sigma). The total protein concentration of lysate was calculated using BCA protein estimation kit (Pierce), as per the manufacturer’s protocols. We used 20 µg of total protein per well, for SDS PAGE. The proteins were separated in 10% poly-acrylamide gel at 110 volts for 2 hours. The proteins were then transferred on PVDF membrane (Millipore, USA) by overnight wet transfer at 30 volts. Immunoblots were blocked in 5% skim milk or 5% BSA for an hour at room temperature. After blocking, the blots were incubated with primary antibodies for IQGAP2 (1: 1000; Abcam), N-cadherin (1: 1000; Abcam), E-cadherin (1: 1000; Abcam), twist1 (1: 1000, Cell Signalling Technology), snail (1: 1000, Cell Signalling Technology), AKT (1: 1000, Cell Signalling Technology), p-AKT473 (1: 1000, Cell Signalling Technology), ERK (1: 1000, Cell Signalling Technology), p-ERK (1: 1000, Cell Signalling Technology) and GAPDH (1: 5000, Sigma) overnight at 4°C. HRP-conjugated, anti-mouse (1:10000, Sigma) and anti-rabbit (1:10000, Sigma) secondary antibodies were then incubated for 1 hour at room temperature followed by detection of bands by chemiluminescence in ChemiDoc XRS+ (Bio-Rad).

### Cell proliferation assay

Cell proliferation assay was performed in 96 well plate using CellTiter 96® AQueous One Solution Reagent (Promega) as per the manufacturer’s protocol. Briefly, 5000 cells/well of from each experimental group (MCF7 IQGAP2_Ex, MCF7 IQGAP2_KD, MDA-MB-468 IQGAP_KD) along with their controls, were plated and incubated for different time intervals. After each time interval, old media was replaced with fresh DMEM (100 µl) mixed with 20 µl of cell titre aqueous solution. The experiment was performed with three technical and three biological replicates. The final absorbance was taken at 495nm between 1- 4 hours. The experiment was performed in triplicate.

### Colony formation assay

Cells from each experimental group (MCF7 IQGAP2_Ex, MCF7 IQGAP2_KD, MDA-MB-468 IQGAP_KD) along with their controls, were seeded into 6-well plates at a density of 1000 cells/well in 10% DMEM and incubated for 10 days at 37°C in 5% CO2. After incubation, the cells were washed with 1X PBS buffer and fixed with methanol + acetic acid (3:1) fixation solution for 5 minutes. Then, cells were washed with 1X PBS buffer and stained with 0.5% crystal violet (MP Biomedicals) for 20 minutes. Cell colonies were photographed by digital camera (Nikon) and were counted using ImageJ software. The experiment was performed in triplicate.

### Wound healing assay

Cells (0.2×10^6^) from each experimental group (MCF7 IQGAP2_Ex, MCF7 IQGAP2_KD, MDA-MB-468 IQGAP2_KD) along with their controls, were seeded in 12-well plates in 10% DMEM. Cells were allowed to grow, till 90-100% confluence. Three parallel lines were drawn with marker on the base of the wells and a perpendicular scratch/wound was created using a 200 µl pipette tip. Cells were cultured for another 24 hours and images were captured using 4X objective lens at different time intervals, i.e. 0, 6, 12 and 24 hours, under an inverted microscope (Nikon). The area of wounds, at 0 and 24 hours for each group, was calculated using ImageJ software. This experiment was conducted in triplicate.

### Transwell cell migration and invasion assay

To perform transwell cell migration assay, we used transwell chambers of 8 μm pore diameter from Millipore. Briefly, 0.05×10^6^ MCF7 cells (IQGAP2_Ex, IQGAP2_KD and their controls) and 0.01×10^6^ MDA-MB-468 cells (IQGAP_KD and it’s control) were seeded in 500 µl of 2% DMEM in the upper chamber of a 12-well plate and 1 ml of 10% DMEM was filled in the lower chamber. The cells were incubated for another 24 hours at 37°C in 5% CO2. After incubation, media was decanted and chambers were washed with 1X PBS buffer. Cells were fixed with 4% PFA for 10 minutes and permeabilised using absolute methanol (Merck) for 20 minutes. Cells were stained with 0.5% crystal violet or Giemsa (HiMedia) for 15 minutes. Cells from the upper chamber were wiped off using a cotton wipe and image was captured using 10X objective lens of an upright bright field microscope (Olympus). Cells from five different fields were counted using NIH ImageJ software. The experiment was performed in triplicates.

For Transwell invasion assay, all steps were same as mentioned in transwell migration assay, except in the initial step, transwell chambers were coated with 100 μl of 1 mg/ml growth factor reduced matrigel (Corning).

### Statistical analyses

Statistical analysis was performed using GraphPad Prism 6.0 Version (GraphPad Software Inc.), Microsoft excel and SPSS. For cell based assays, continuous data was presented as the mean ± standard error of the mean. Student’s t-test (2-tailed, unpaired) was used to determine significant difference between the groups. For IHC data, significance of difference in distribution frequency of Allred scores, between tumor and control group, was determined using 2-tailed, chi-square test. Spearman correlation test was also used to determine correlation between clinicopathological parameters and Allred scores for IQGAP2 expression. P-value ≤ 0.05, was considered to be significant for all the tests.

## Results

### IQGAP2 expression doesn’t correlate with breast cancer molecular subtype

To determine the correlation of molecular subtypes of breast cancer with IQGAP2 expression we used cells (MCF7, T-47D, MCF 10A, MDA-MB-453, MDA-MB-468 and MDA-MB-231) representing different subtypes, and determined mRNA and protein expression levels. We observed very low expression of IQGAP2 protein in T-47D, MCF 10A and MDA-MB-231 breast cancer cell lines and higher expression in MCF7, MDA-MB-453, MDA-MB-468 cell lines (Figure 1A). Similar observation was made at mRNA level (Figure 1B). Cells in higher expression and lower expression groups are mixed population, containing ER/PR positive and triple negative cell lines. Therefore, IQGAP2 expression cannot be correlated with breast cancer molecular subtype.

**Figure 1.**
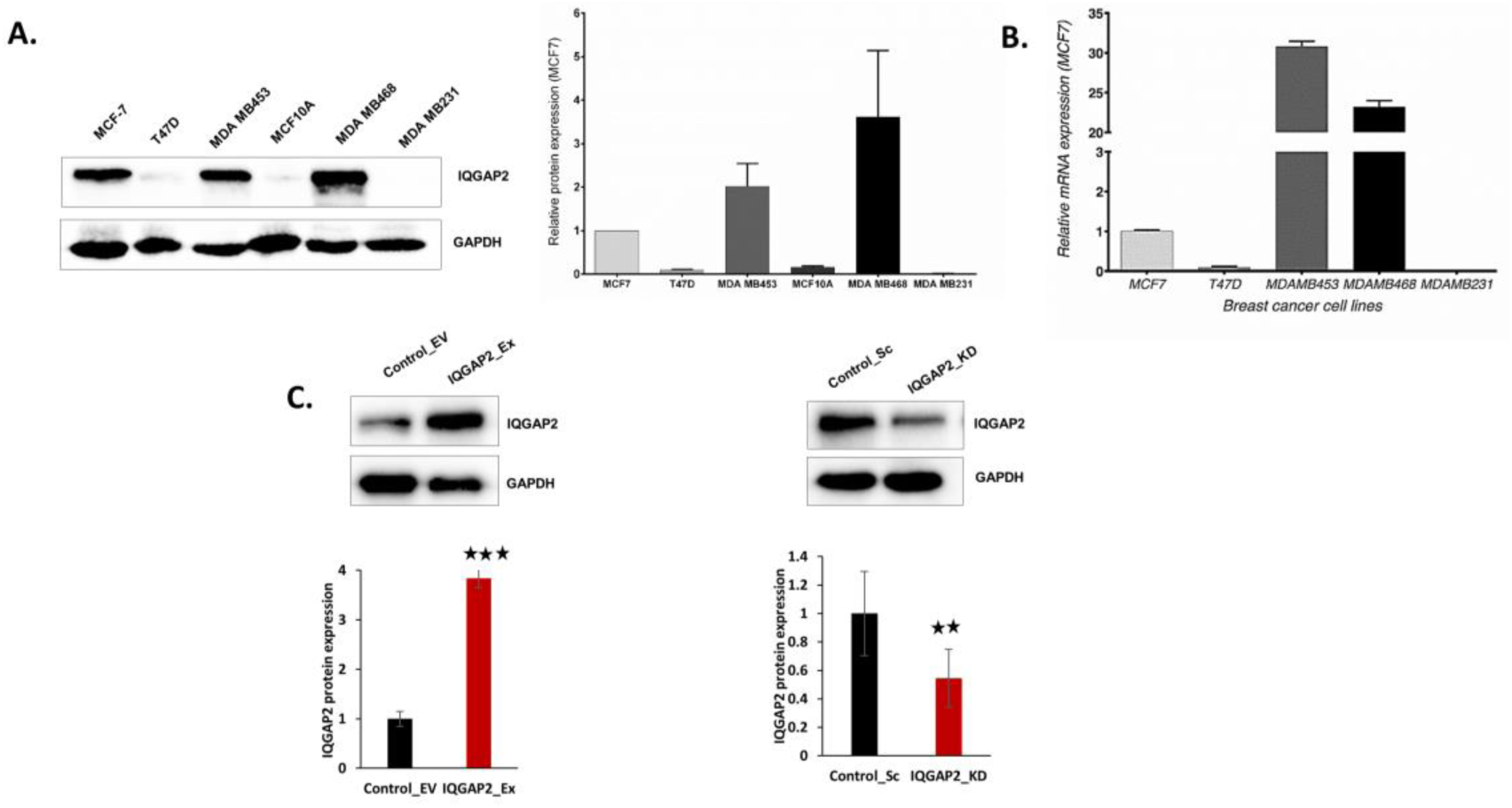
Expression levels of IQGAP2 in different breast cancer cell lines. A) The western blot of IQGAP2 showing its endogenous expression in different breast cancer cell lines of specific molecular signature (Left panel). Here, IQGAP2 expression is shown in MCF7 (ER/PR positive), T-47D (ER/PR/Her2 positive), MDA-MB-453 (Her2 positive), MCF 10A (ER/PR/Her2 negative normal like), MDA-MB-468 (ER/PR/Her2 negative) and MDA-MB-231 (ER/PR/Her2 negative and highly aggressive) cell lines. Right panel shows densitometry bar graph (relative to the expression of MCF7) of the western blot. (N = 3). B) The endogenous mRNA level of IQGAP2 (relative to the endogenous expression in MCF7) in different breast cancer cell lines. (N = 3). C) The upper panels show representative western blot images of stable lines of MCF7 Control_EV, MCF7 IQGAP2_Ex, MCF7 Control_Sc and, MCF7 IQGAP2_KD, respectively and the lower panels showing the relative densitometry bar graph of respective images calculated with ImageJ software. The graphs show a significant change in over expression (p ≤ 0.001) and knockdown (p ≤ 0.01) of IQGAP2 in MCF7 IQGAP2_Ex and MCF7 IQGAP2_KD, respectively, compared to their controls. (N=3, 2-tailed unpaired t-test). * represents *p* value ≤ 0.05, ** represents *p* value ≤ 0.01, *** represents *p* value ≤ 0.001, N represents experiment replicates.

### Reduced IQGAP2 expression promotes cell proliferation in ER/PR positive cells

Further, we used estrogen receptor positive cell line, MCF7 to check the effect of IQGAP2 expression levels on the rate of proliferation, which is one of the hallmarks of tumorigenesis. We used MCF7 cells with stable ectopic expression and depletion of IQGAP2 (Figure 1C).

In MTT based cell proliferation assay, we observed that the MCF7 with IQGAP2 depletion were proliferating at significantly (p ≤ 0.0001) higher rate than the control group (Cell number, IQGAP2_KD/ Control_Sc = 47065±5335/19937±1173, post 96 hours of cell plating) (Figure 2A) whereas an opposite trend was observed in MCF7 cells with ectopic expression of IQGAP2 (Cell number, IQGAP2_Ex/ Control_ EV = 40288±441/ 42633±349, post 96 hours of cell plating) (Figure 2B). In colony formation assay similar trend was observed. In MCF7 with IQGAP2 depletion, more colonies were observed compared to control group (96±13 versus 37±12). The size of the colonies was also measured and significantly (p ≤ 0.0001) larger colonies were observed in MCF7 with reduced IQGAP2 expression, compared to the controls (Figure 2C). We observed the opposite trend in MCF7 with IQGAP2 ectopic expression, where significantly (p ≤ 0.001) less number (4.7±0.6 versus 48±5) and smaller colonies were observed, compared to the control group (Figure 2D). These results support the role of IQGAP2 as tumor suppressor gene.

**Figure 2.**
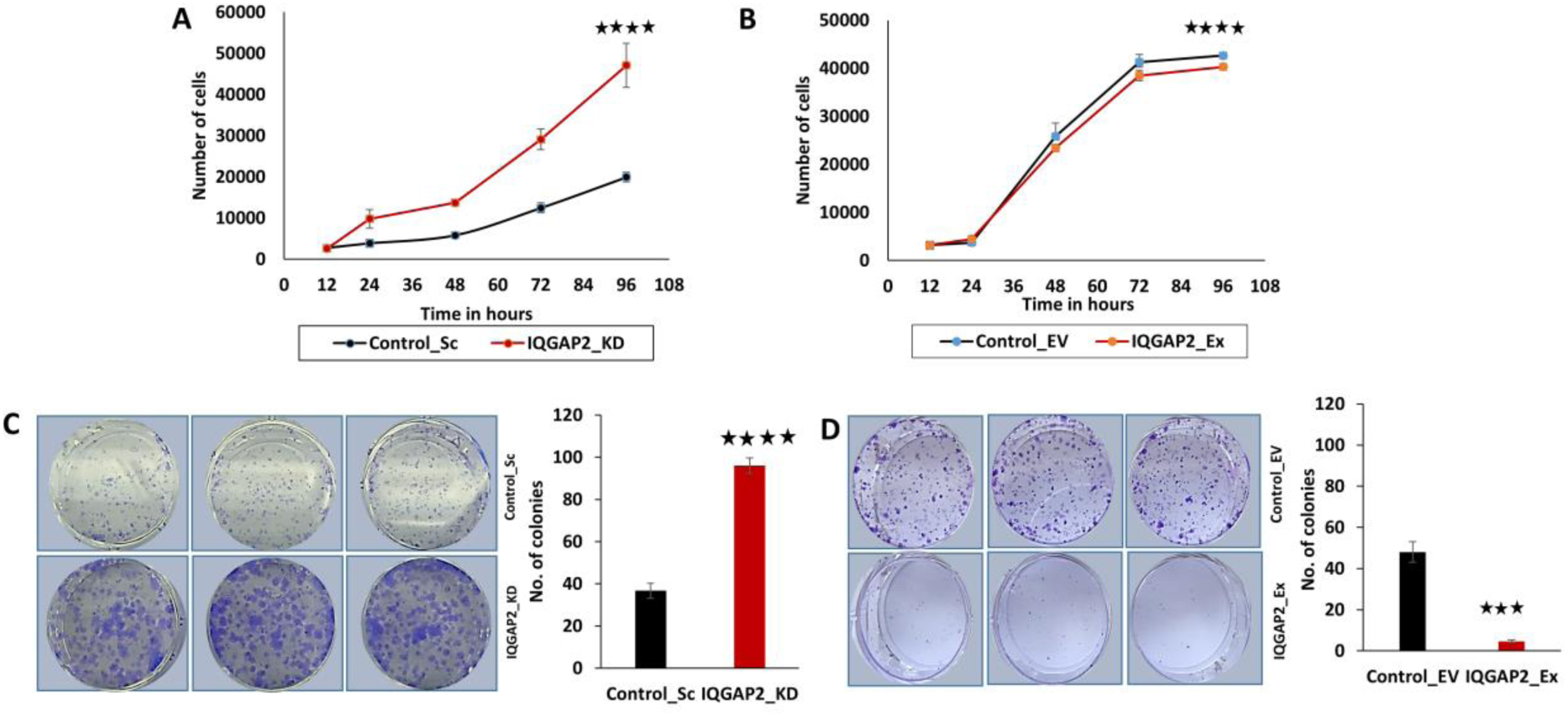
Reduced IQGAP2 expression promotes cell proliferation and cell growth in breast cancer cell lines. A) The graph shows comparisons between cell number of MCF7 IQGAP2_ KD group and MCF7 Control_Sc group, at different time intervals. Here, y-axis indicates the cell number and x-axis shows time in hours. The cell number was calculated using MTT assay. Knockdown of IQGAP2 shows significant increase in cell proliferation (*N* = 3, 2-tailed unpaired t-test, *p* value ≤ 0.0001). B) The graph represents result of MTT assay, showing the cell number of MCF7 IQGAP2_Ex group and MCF7 Control_EV group. Over expression of IQGAP2 shows significant decrease in cell proliferation (*N* = 3, 2-tailed unpaired t-test, *p* value ≤ 0.0001). C) Left panel shows the images of colony formation assay in MCF7 IQGAP2_KD group and MCF7 Control_Sc group. The bar graph in the right panel shows the differences in colony numbers between both the groups. Knockdown of IQGAP2 shows significantly more number of colonies (*N* = 3, 2-tailed unpaired t-test, *p* value ≤ 0.0001). D) Image showing the number of colonies in MCF7 IQGAP2_Ex group and MCF7 Control_EV group (left panel). The right panel illustrations bar graph, showing the differences of colony numbers between both the groups. The colony number was calculated using ImageJ software. Over expression of IQGAP2 shows significantly low colony number (*N* = 3, 2-tailed unpaired t-test, *p* value ≤ 0.001), compared to control. * represents *p* value ≤ 0.05, ** represents *p* value ≤ 0.01, *** represents *p* value ≤ 0.001, **** represents *p* value ≤ 0.0001, N represents experiment replicates.

### IQGAP2 silencing promotes migration and invasion in ER/PR positive breast cancer cells and vice versa

The cell migratory property, which is another important property of cancer, was assessed using wound healing and transwell migration assay. As shown in figure 3, inhibition of IQGAP2 expression in MCF7 cells, resulted in increased rate of wound healing as well as in transwell migration. In wound healing assay, 92% of the wound was recovered after 24 hours of wound formation, in MCF7 cells with IQGAP2 knockdown, whereas only 75% of the wound recovery was recorded in control group (Figure 3A), which is significantly (p ≤ 0.0001) higher. Transwell migration assay showed that significantly (p ≤ 0.0001) more MCF7 cells migrated through the membrane in IQGAP2 knockdown group (128.6±13.75 cells/field), compared to the control group (50.2±4.54 cells/field) (Figure 3C).

**Figure 3.**
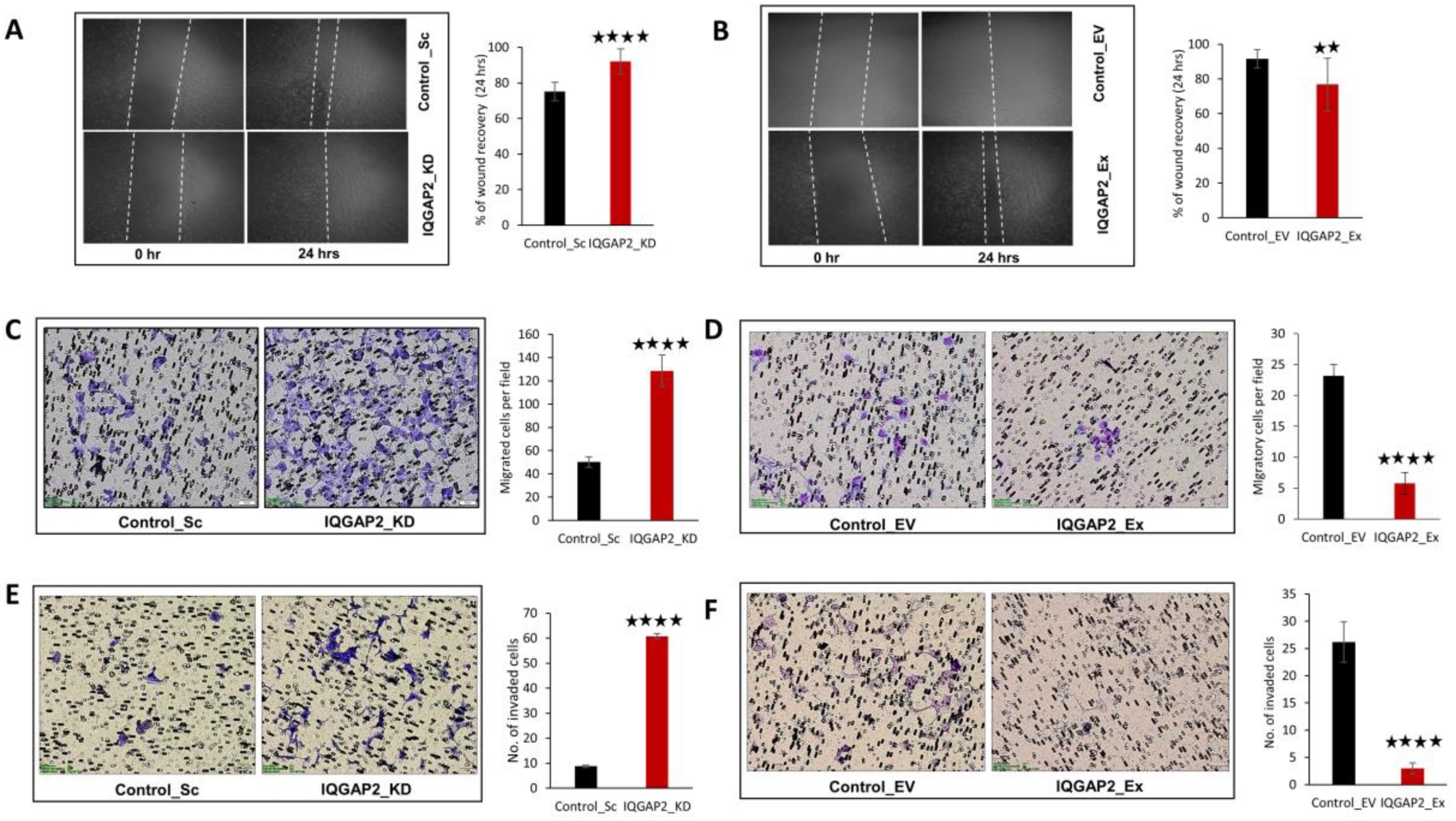
IQGAP2 expression level affects migration and invasion of ER/PR positive breast cancer cells. **A)** Representative images of wound healing assay in MCF7 IQGAP2_KD and MCF7 Control_Sc group, at 0 hour and 24 hours (left panel). The bar graph (right panel) shows percentage recovery of wound after 24 hours. Here, y-axis represents the percentage of wound recovery after 24 hours and x-axis shows type of cells used for assay. Graph shows higher wound recovery rate of IQGAP2 knockdown group (92%), compared to empty vector control (75%) (*N* = 3, 2-tailed unpaired t-test, *p* value ≤ 0.0001). **B)** Left panel shows representative images of wound healing assay of MCF7 IQGAP2_Ex and MCF7 Control_EV cells at 0 and 24 hours. Bar graph in right panel shows the percentage recovery of wound post 24 hours of wound formation. Here, y-axis represents the percentage of wound recovery after 24 hours and x-axis shows type of cells used for assay. Ectopic expression of IQGAP2, shows slow wound recovery (76.7%) compared to vector control (91.6%) (*N* = 3, 2-tailed unpaired t-test, *p* value ≤ 0.01). **C)** Representative images (left panel) of transwell migration assay in MCF7 IQGAP2_KD and MCF7 Control_Sc group, showing higher migratory properties of IQGAP2_KD group. The bar graph (right panel) shows the number of migrated cells in both the groups post 24 hours of cell plating. Here, y-axis represents the number of cells migrated in 24 hours and x-axis shows type of cells used for assay. MCF7 cells with IQGAP2 knockdown show enhanced (128.6±13.75 cells/field) transwell migration, compared to scrambled vector (50.2±4.54 cells/field) (*N* = 3, 2-tailed unpaired t-test, *p* value ≤ 0.0001). **D)** Representative images (left panel) of transwell migration assay in MCF7 IQGAP2_Ex and MCF7 Control_EV cells. The bar graph (right panel) shows the number of migrated cells in both the groups post 24 hours of cell plating. Here, y-axis represents the number of cells migrated in 24 hours and x-axis shows type of cells used for assay. MCF7 cells with ectopic IQGAP2 expression show reduced (5.75±2.62 cells/field) transwell migration, compared to control vector (23.2±1.78 cells/field) (*N* = 3, 2-tailed unpaired t-test, *p* value ≤ 0.0001). **E)** Representative images (left panel) of transwell invasion assay in MCF7 IQGAP2_KD and MCF7 Control_Sc group, showing more invasive properties of IQGAP2_KD group The bar graph (right panel) shows the number of migrated cells in both the groups post 24 hours of cell plating. Here, y-axis represents the number of cells migrated in 24 hours and x-axis shows type of cells used for assay. MCF7 cells with IQGAP2 knockdown increase (60.66±1.15 cells/field) transwell invasion, compared to scrambled vector 8.75±0.5 cells/field) (*N* = 3, 2-tailed unpaired t-test, *p* value ≤ 0.0001). **F)** Representative images of transwell invasion assay in MCF7 IQGAP2_Ex and MCF7 Control_EV cells. The bar graph (right panel) shows the number of migrated cells in both the groups post 24 hours of cell plating. Here, y-axis represents the number of cells migrated in 24 hours and x-axis shows type of cells used for assay. MCF7 cells with ectopic IQGAP2 expression showing reduced (3±1 cells/field) transwell invasion, compared to control vector (26.2±3.7 cells/field) (*N* = 3, 2-tailed unpaired t-test, *p* value ≤ 0.0001). * represents *p* value ≤ 0.05, ** represents *p* value ≤ 0.01, *** represents *p* value ≤ 0.001, **** represents *p* value ≤ 0.0001, N represents experiment replicates. Scale bar in all images is 50 microns.

In order to check effect of IQGAP2 on cell invasiveness, transwell chamber assay was performed with Matrigel. In IQGAP2 knockdown condition significantly (p ≤ 0.0001) more MCF7 cells (60.66±1.15 cells/field) invaded through the Matrigel layer and moved to the bottom surface of the chamber, compared to the control group (only 8.75±0.5 cells/field) (Figure 3E).

To further investigate the effects of IQGAP2 on breast cancer cells, we used MCF7 cells with stable overexpression of IQGAP2 (IQGAP2_Ex) and control (Control_EV). Ectopic IQGAP2 expression significantly decreased cell migration and invasion. In Wound healing assay, 91.6% of the wound was recovered in control group whereas only 76.7% of wound was recovered in IQGAP2 ectopic expressed group (p ≤ 0.01) (Figure 3B). Similar trend was observed in transwell chamber assay, where very less (5.75±2.62 cells/field) MCF7 cells, from ectopic IQGAP2 expression group, migrated through chamber compared to the control group (23.2±1.78 cells/field) (p ≤ 0.0001) (Figure 3D). In cell invasion assay, we observed fewer cells (3±1 cells/field) invaded through chamber, in IQGAP2 ectopic expression group, compared to the control (26.2±3.7 cells/field) (p ≤ 0.0001) (Figure 3F). The results indicate that overexpression of IQGAP2 in MCF7 cells led to significant reduction of cell migration and breast cancer metastasis in vitro.

### Effect of reduced IQGAP2 expression on tumorigenic cell properties is irrespective of ER/PR status

To find out if the effect of IQGAP2 on cell proliferation, migration and invasion, is irrespective of molecular subtype of breast cancer, we used MDA-MB-468 cells, which are triple negative. In MDA-MB-468 cell lines the endogenous level of IQGAP2 was very high (Figure 1A) hence we depleted IQGAP2 in these cells stably (Figure 4A).

**Figure 4.**
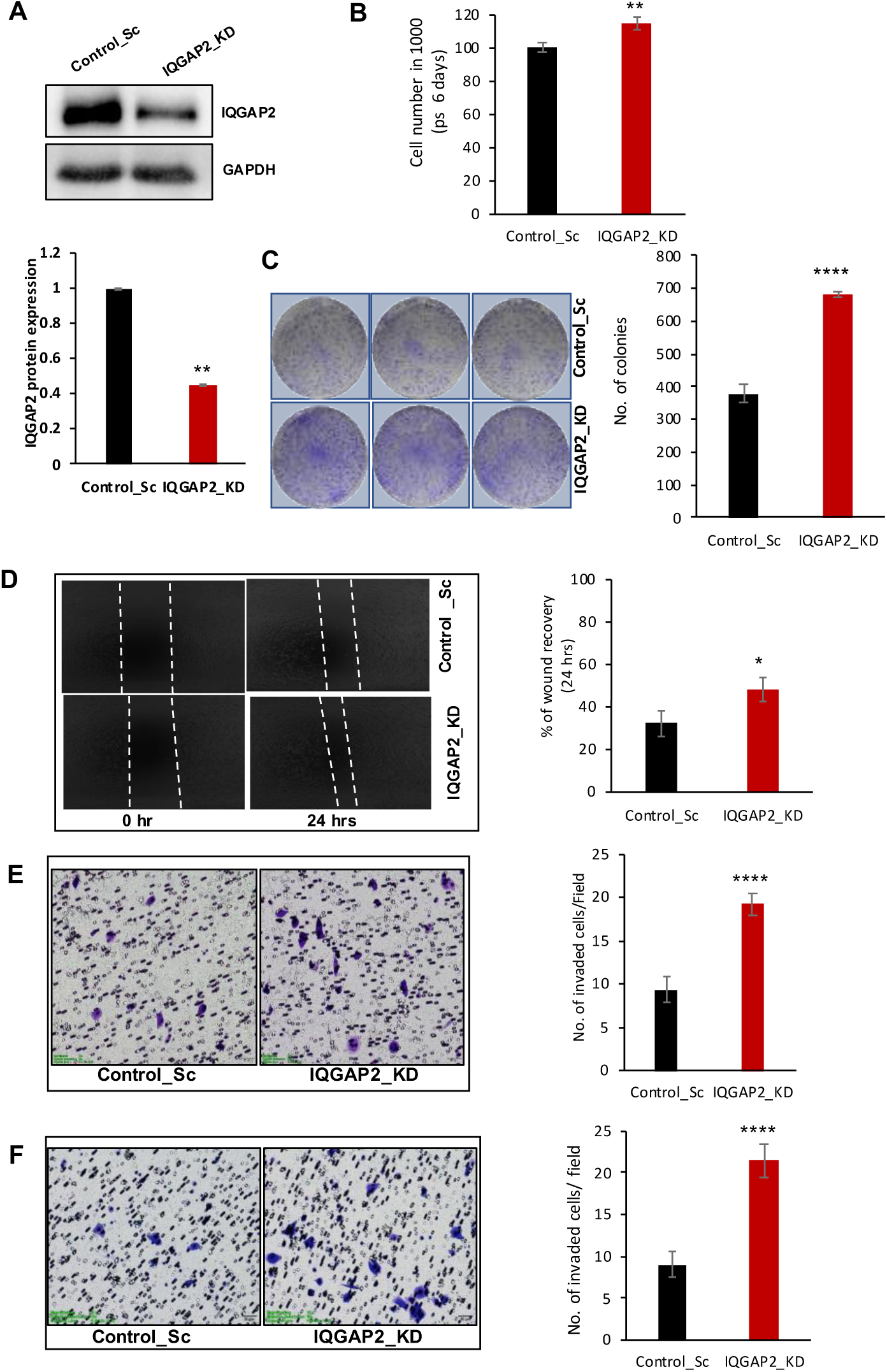
Reduced expression of IQGAP2 promotes tumorigenic properties in triple negative breast cancer cell line. **A)** Western blot image showing (upper panel) knock down of IQGAP2 in MDA-MB-468 cell lines and its relative densitometry bar graph (lower panels). The graph shows a significant decrease in IQGAP2 expression level in knockdown stable cell line. (N=3, 2-tailed unpaired t-test, p ≤ 0.01). **B)** Bar graph shows the comparisons between cell number in MDA-MB-468 IQGAP2_KD group and Control_Sc group, post 5 days of cell plating. MTT assay was performed to calculate the cell number. Knockdown of IQGAP2 shows higher cell proliferation rate (*N* = 3, 2-tailed unpaired t-test, p value ≤ 0.01), compared to control. **C)** Representative images of colony formation assay (left panel) in MDA-MB-468 IQGAP2_KD group and Control_Sc group. In the right panel bar graph shows the difference of colony numbers between both the groups. Knockdown of IQGAP2 shows significantly more number of colonies (*N* = 3, 2-tailed unpaired t-test, *p* value ≤ 0.0001), compared to control. **D)** Left panel shows representative images of wound healing assay of MDA-MB-468 IQGAP2_KD and Control_Sc cells at 0 and 24 hours. Bar graph in right panel shows the percentage recovery of wound post 24 hours of wound formation. Here, y-axis represents the percentage of wound recovery after 24 hours and x-axis shows type of cells used for assay. Graph shows higher wound recovery rate of IQGAP2 knockdown group (48.45±5.31%), compared to empty vector control (32.69±5.96%) (*N* = 3, 2-tailed unpaired t-test, *p* value ≤ 0.05). **E)** Representative images (left panel) of transwell migration assay in MDA-MB-468 IQGAP2_KD and Control_Sc group, showing higher migratory properties of IQGAP2_KD group. The bar graph (right panel) shows the number of migrated cells in both the groups post 24 hours of cell plating. Here, y-axis represents the number of cells migrated in 24 hours and x-axis shows type of cells used for assay. MDA-MB-468 cells with IQGAP2 knockdown show enhanced transwell migration (19.2±1.3 cells/field), compared to control (9.4±1.5 cells/field) (*N* = 5, 2-tailed unpaired t-test, *p* value ≤ 0.0001). **F)** Representative images (left panel) of transwell invasion assay in MDA-MB-468 IQGAP2_KD and Control_Sc group, showing more invasive properties of IQGAP2_KD group. The bar graph (right panel) shows the number of migrated cells in both the groups post 24 hours of cell plating. Here, y-axis represents the number of cells migrated in 24 hours and x-axis shows type of cells used for assay. MDA-MB-468 cells with IQGAP2 knockdown show enhanced transwell invasion (21.4±2.07 cells/field), compared to control (9±1.58 cells/field) (*N* = 5, 2-tailed unpaired t-test, *p* value ≤ 0.0001). * represents *p* value ≤ 0.05, ** represents *p* value ≤ 0.01, *** represents *p* value ≤ 0.001, **** represents *p* value ≤ 0.0001, N represents experiment replicates. Scale bar in all images is 50 microns.

In MDA-MB-468 cell lines, a reduced IQGAP2 expression showed significantly (p ≤ 0.01) higher cell proliferation rate compared to the control group (Figure 4B). In colony formation assay, significantly (p ≤ 0.0001) more number of colonies were observed in IQGAP2 knock down group (680.3±10.2 cells/well) compared to the control group (377±28.3 cells/well) (Figure 4C). Similarly, in MDA-MB-468, reduction in IQGAP2 level resulted in increased migration and invasion of cells. In wound healing assay, significantly (p ≤ 0.05) more wound recovery was noted in IQGAP2 knockdown group (48.45±5.31%) compared to the control (32.69±5.96%) (Figure 4D). Transwell migration assay was also performed with MDA-MB-468 cells. This assay showed significantly (p ≤ 0.0001) more invaded cells towards bottom of the transwell in IQGAP2 knockdown group (19.2±1.3 cells/field) than the control (9.4±1.5 cells/field) group (Figure 4E). Invasion assay also showed same trend as migration assay. Here, significantly (p ≤ 0.0001) more number of cells were observed toward the lower chamber of the insert in IQGAP2 knockdown group (21.4±2.07 cells/field) than the control group (9±1.58 cells/field) (Figure 4F). All of these results suggest the tumorigenic effect of IQGAP2 reduction is irrespective of molecular subtype of breast cancer cell.

### IQGAP2 induces epithelial-mesenchymal transition (EMT) in breast cancer cells

To determine the underlying mechanisms of IQGAP2’s role in tumorigenesis, we tested the effect of it’s expression level on EMT; western blot analysis was performed to detect the expression of epithelial and mesenchymal protein markers. Our results showed that over expression of IQGAP2 reduces the expression of mesenchymal marker proteins: twist, and N-cadherin while it up-regulated epithelial marker, E-cadherin (Figure 5A). Conversely, MCF7 cells with reduced IQGAP2 levels, exhibited decreased E-cadherin expression, but increased expression of EMT markers, twist and N-cadherin. (Figure 5B). Overall, these results suggest that increased expression of IQGAP2 can inhibit tumor invasion and metastasis by suppressing EMT.

**Figure 5.**
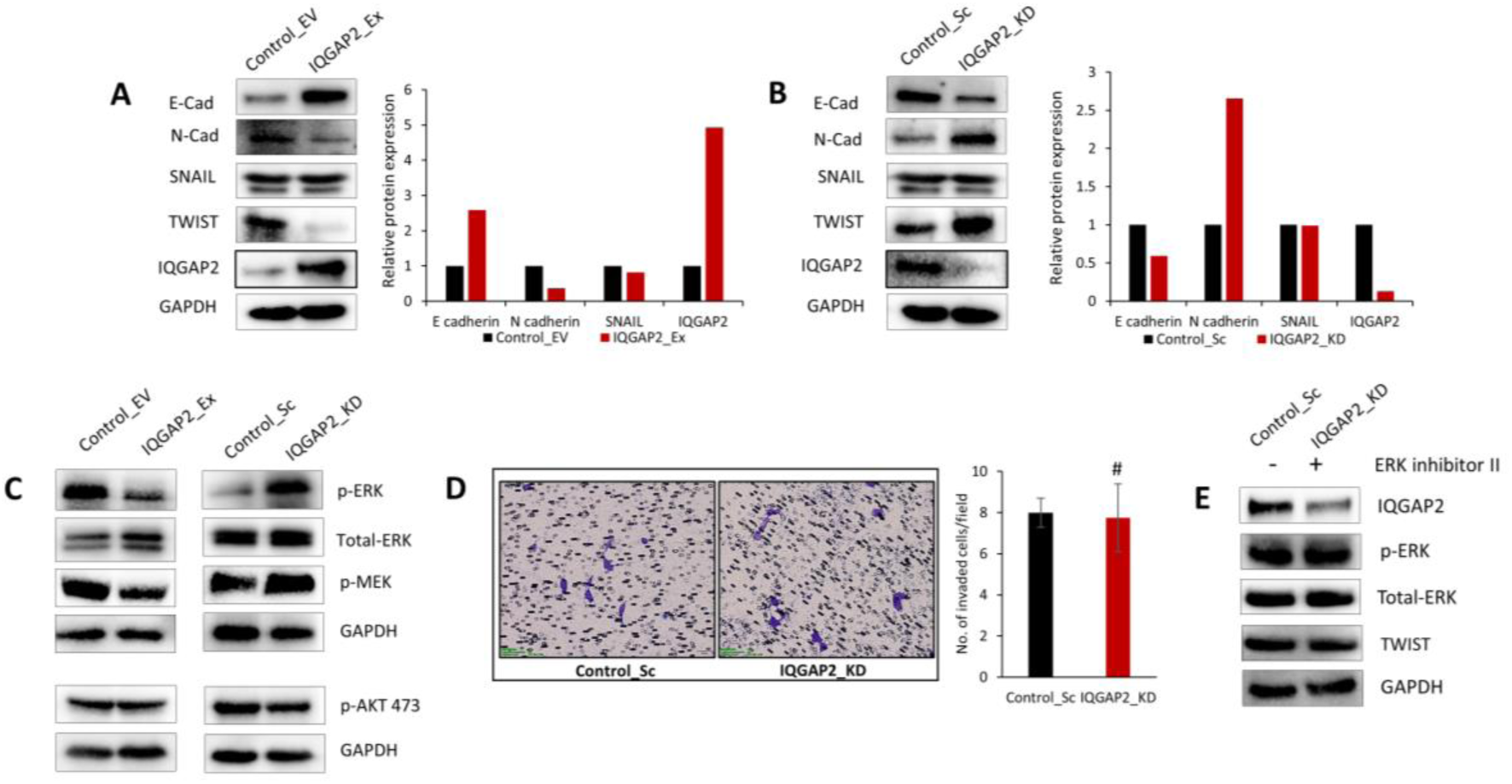
Reduced IQGAP2 expression induces epithelial to mesenchymal transition in MCF7 cells via the MEK/ERK signalling pathways. **A)** Western blot image showing expression levels of different EMT markers and loading control (GAPDH) in MCF7 IQGAP2_KD and Control_Sc groups. In right panel bar graphs shows the relative densitometry of EMT markers. **B)** Western blot image of different EMT markers in MCF7 IQGAP2_Ex and Control_EV groups. In right panel, bar graph shows the relative densitometry of EMT markers. **C)** Western blot image of major pathways associated with IQGAP2 in MCF7. Here in right panel ERK and MEK show more phosphorylation with IQGAP2 reduction, than control but there was no change in AKT. Just opposite pattern is observed in left panel (MCF7 IQGAP2_Ex and Control_EV) for ERK; there also, AKT activation is not affected. **D)** Image of invasion assay. Left panel shows MCF7 IQGAP2_KD cells treated with ERK inhibitor (U0126) 10 µM, right panel shows MCF7 Control_Sc (treated with solvent). The bar graph shows the number of cells invaded matrigel coated cell chamber at 24 hours from MCF7 IQGAP2_KD cells treated with ERK inhibitor (U0126) and from MCF7 Control_Sc (treated with solvent). Both the groups show no statistically significant difference (*N* = 3, 2-tailed unpaired t-test, *p* value >0.05). **E)** Western blot analysis of different EMT markers after ERK inhibition in IQGAP2 knock down MCF7 cell lines. Right lane shows MCF7 IQGAP2_KD cells treated with ERK inhibitor (U0126) 10 µM. Left lane shows MCF7 Control_Sc (treated with solvent). Both the lanes show similar expression levels of Twist. E-cad and N-cad stand for E-Cadherin and N-Cadherin, respectively. N represents experiment replicates, # represents non-significant at p ≤ 0.05. Scale bar in all images is 50 microns.

### IQGAP2 mediated EMT occurs via the MEK/ERK signalling pathways in MCF7 cells

MEK/ERK and PI3K/AKT signalling pathways play an important role in migration, invasion, and metastasis of cancer by regulating EMT; therefore, we investigated if IQGAP2 was capable of promoting EMT via the MEK/ERK and PI3K/AKT pathways in MCF7 cells. Firstly, we estimated the alteration of ERK and AKT. As shown in figure 5C, overexpression of IQGAP2, decreased phosphorylation of ERK but not AKT, while silencing of IQGAP2 expression, increased ERK phosphorylation but did not show a change in AKT phosphorylation. This data suggested that reduction in IQGAP2 was promoting EMT via MEK/ERK activation.

Further, to evaluate whether the effects of IQGAP2 on cell migration and invasion were dependent on the ERK pathway, MCF7 line with stable IQGAP2 knock down, was treated with ERK inhibitor (U0126) (Calbiochem) 10 µM for 30 min. Inhibition of ERK abrogated the higher invasion observed with IQGAP2 knock down, and there was no significant difference compared to control (Figure 5D). In addition, inhibition of ERK1/2 by U0126 reversed the effect on twist expression, that is up-regulation which was induced by IQGAP2 knockdown (Figure 5E). These results confirm that reduced IQGAP2 expression induces cell migration, invasion, and EMT through MEK/ERK signalling pathways.

### IQGAP2 is down-regulated in human breast cancer patients

To validate our findings at cell line level, we used breast cancer patient tissue samples, to check the expression levels of IQGAP2 in tumor tissue compared to normal tissue. Here, we analysed the expression pattern and localization of IQGAP2 at protein level in surgically resected breast tumor tissues and their adjacent normal (uninvolved) region of 195 breast cancer patient, using IHC. Comparison of Allred score for IQGAP2 in tumor and normal tissue, showed significant (p ≤ 0.0001) reduction of IQGAP2 expression in tumor tissue (median value =3, range = 0 - 8) compared to normal tissue (median value =8, range = 5 - 8) (Figure 6A). In terms of percentage distribution of IQGAP2 expression, most of the normal region were positive for strong (79.36%, 50 out of 63) and moderate staining (20.63%, 13 out of 63) wherein staining in tumor area was weak to negative in 37.43% (73 out of 195), moderate in 49.23% (96 out of 195) and strong only in 13.33% cases (26 out of 195) (Figure 6B). Further, we checked the localization of IQGAP2, in normal breast tissue and tumor tissue. The IQGAP2 expression was predominantly observed in cytosolic region of the cells in normal as well as in tumor tissue (Figure 6C).

**Figure 6.**
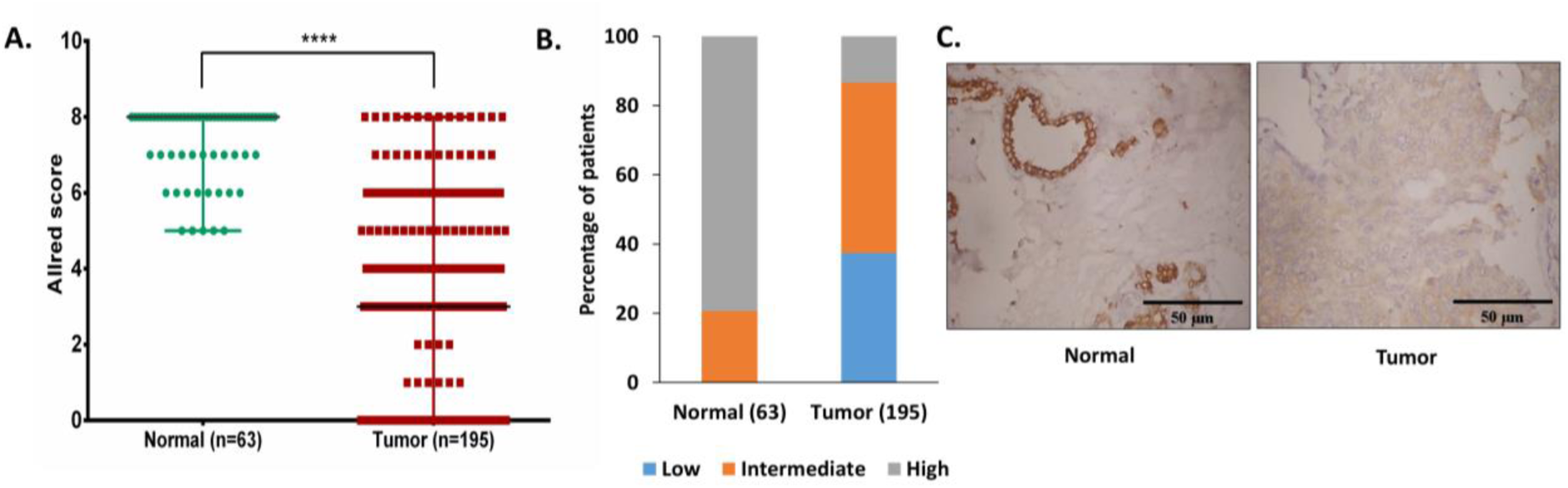
IQGAP2 is down regulated in tumor tissues of breast cancer patients. A) The comparison of Allred scores of IQGAP2 expression between tumor (n = 195) and adjacent normal tissue (n = 63) of breast cancer patients. The Allred score for IQGAP2 is significantly (p ≤ 0.0001, Mann-Whitney U test) reduced in tumor tissues compared to normal. B) The percentage frequency distribution of normal vs cancer tissues according to the Allred score of IQGAP2. The tissues were grouped into three; Low (AS = 0-2), intermediate (AS = 3-6) and high (AS = 7-8). y-axis represents percentage of patient positive for particular IQGAP2 staining Allred score. x-axis represents normal and tumor tissue groups. C) The representative images of tumor tissue and adjacent normal tissue of breast cancer patient, showing IQGAP2 expression and localization. The images were captured at 10X objective lens of bright field microscope. Scale bar in all images is 50 microns. n = 3, AS=Allred Score, **** represents *p* value ≤ 0.0001.

To find out the correlation between IQGAP2 expression and clinic-pathological features, we divided patients into two groups, IQGAP2 negative and IQGAP2 positive based on IQGAP2 IHC score (Allred score 0-2- negative, Allred score 3-8- positive). Chi-square test was performed to analyse the correlation between these groups for different clinic-pathological characteristics (age, tumor size, lymph node metastasis, lymphovascular invasion and histological grade). Analysis showed that reduced IQGAP2 expression in breast cancer was significantly correlated with age (p ≤ 0.05), lymphovascular invasion (p ≤ 0.001) and histological grade (p ≤ 0.05) but not with lymph node metastasis and size of the tumor of patient (Table 2).

**Table 2.**
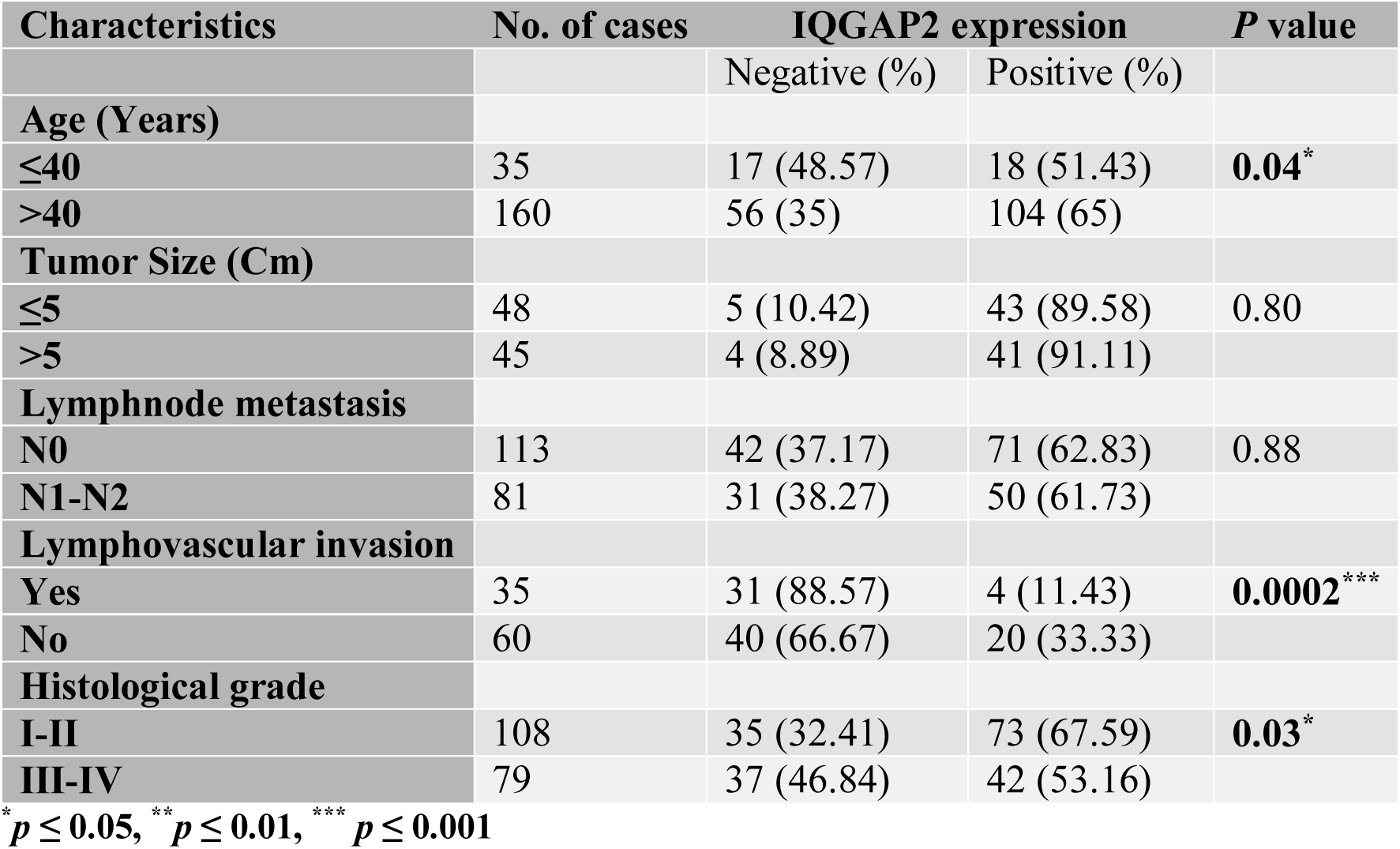
Correlation of IQGAP2 expression with histopathological parameters of breast cancer.

| Characteristics | No. of cases | IQGAP2 expression |  | *P* value |
| --- | --- | --- | --- | --- |
|  |  | Negative (%) | Positive (%) |  |
| **Age (Years)** |  |  |  |  |
| ≤40 | 35 | 17 (48.57) | 18 (51.43) | **0.04\*** |
| >40 | 160 | 56 (35) | 104 (65) |  |
| **Tumor Size (Cm)** |  |  |  |  |
| ≤5 | 48 | 5 (10.42) | 43 (89.58) | 0.80 |
| >5 | 45 | 4 (8.89) | 41 (91.11) |  |
| **Lymphnode metastasis** |  |  |  |  |
| N0 | 113 | 42 (37.17) | 71 (62.83) | 0.88 |
| N1-N2 | 81 | 31 (38.27) | 50 (61.73) |  |
| **Lymphovascular invasion** |  |  |  |  |
| Yes | 35 | 31 (88.57) | 4 (11.43) | **0.0002\*\*\*** |
| No | 60 | 40 (66.67) | 20 (33.33) |  |
| **Histological grade** |  |  |  |  |
| I-II | 108 | 35 (32.41) | 73 (67.59) | **0.03\*** |
| III-IV | 79 | 37 (46.84) | 42 (53.16) |  |

\* $p \leq 0.05$ , \*\* $p \leq 0.01$ , \*\*\* $p \leq 0.001$

## Discussion

Breast cancer is the second driving reason of cancer deaths among women. In few decades, the advancement in surgery, chemotherapy and targeted therapy have improved the survival of the patient but long-term survival of the patient is still a big challenge, because of the drug resistance and relapse of the disease. Identification of new molecular markers in breast cancer will help in early diagnosis and cure against drug resistance of this disease and will improve the survival of the patient. In this study, we have explored the role and signalling mechanism of IQGAP2 scaffolding protein in breast cancer progression. Currently, the role and detailed signalling mechanism of IQGAP2 in many cancers including breast cancer is not yet explored and needs immediate attention so that new possibilities of therapy against cancer could be achieved.

In the current study, our immunohistochemistry results have shown that IQGAP2 is significantly reduced in a large proportion of breast cancer cases which is consistent with the RNA data analysed from Oncomine and TCGA database [22]. Invasion of a cancer to the proximate blood vessels and/or lymphatic system i.e. Lymphovascular invasion, is one of the leading event for metastasis and it is correlated with poor survivability of the patient [24] [25]. Here, we have observed that more patients with IQGAP2 negative or low expression were positive for lymphovascular invasion. According to AJCC TNM stage and grading system higher stage i.e. III and IV represents the highly invasive and metastasis condition of a cancer patient [26]. We have found that the negative or weak IQGAP2 expression had a higher frequency in patients with higher stages, III or IV, compared to patients with early stages of breast cancer and vice versa. Remarkably, we did not find a significant correlation among IQGAP2 expression and tumor size. Here, we suggest that the change in late stages of IQGAP2 affect migration and invasion (metastasis) more, rather than cell growth, which starts in the early stages of cancer. We also find a significant correlation of age with IQGAP2 expression; there was higher frequency of IQGAP2 negative patients in ≤40 year group, suggesting that no/low expression of IQGAP2 exposes to cancer early. On the basis of the above correlation studies of IQGAP2 in breast cancer and the survival data published recently by our group [22], present study suggests that IQGAP2 has a protective role and it functions as a tumor suppressor in breast cancer patients.

To validate the function of IQGAP2 on in-vitro breast cancer model, different cell lines representing specific molecular sub-type of breast cancer were investigated. Initially, endogenous expression of this gene at protein and RNA level was measured that did not show any specificity towards molecular sub-type. The ER/PR positive MCF7 cell lines on reduction of IQGAP2 level show increase tumorigenic properties i.e. increased cell proliferation, migration and invasion. The opposite trend was observed with ectopic expression of IQGAP2. Two major pathways, ERK and AKT promote migration and invasion of ER/PR positive breast cancer [27]. Previous study has also shown the role IQGAP2 in AKT pathways in prostate cancer [17]. In our study knockdown of IQGAP2 in MCF7 activated MEK-ERK signalling pathways. Interestingly, we did not observe a change in AKT pathway. Earlier reports suggest that activation of MEK-ERK cascade leads to downregulation of E-cadherin [28] and upregulation of EMT markers, N-cadherin, twist, snail and slug [29]. In our study, we also observed the same trend. Reduction of IQGAP2, increases the expression levels of N-cadherin, twist and slug and reduces E- cadherin level. This effect gets rescue after ERK/MEK inhibitor treatment in IQGAP2_KD group. An opposite pattern was also observed when IQGAP2 ectopically expressed in MCF7.

We further checked functions of IQGAP2 in triple negative breast cancer cell line, MDA-MB-468. A similar tumorigenic property of IQGAP2 reduction was observed as MCF7. The reduction in IQGAP2 level increases the cell growth, migration and invasion properties of MDA-MB-468. Altogether, IQGAP2 reduction promotes cancer progression and metastasis irrespective of the molecular subtypes of breast cancer.

In conclusion, our study explored, for the first time, the overall expression pattern of IQGAP2 in breast cancer patients, the association of IQGAP2 with the histological parameters of breast cancer and the detailed mechanism behind IQGAP2 function in breast cancer development and progression. Our data indicates that reduced IQGAP2 expression promotes cell proliferation migration and invasion via. activation of ERK pathway and plays a crucial role in the development of breast cancer and hence it may be a potential therapeutic target in breast cancer.

